# P4HA2 is associated with prognosis, promotes proliferation, invasion, migration and EMT in glioma

**DOI:** 10.1101/2020.02.05.935221

**Authors:** Jing Lin, Xiao-Jun Wu, Wen-Xin Wei, Xing-Chun Gao, Ming-Zhu Jin, Yong Cui, Wei-Lin Jin, Guan-Zhong Qiu

**Author notes:** Contributed equally (Jing Lin, Xiao-Jun Wu, Wen-Xin Wei). Corresponding author: Guanzhong Qiu,; Weilin Jin, E-mail: or.

## Abstract

Prolyl-4-hydroxylase subunit 2 (P4HA2), as a member of collagen modification enzymes, is induced under hypoxic conditions with essential roles in the collagen maturation, deposition as well as the remodeling of extracellular matrix(ECM). Mounting evidence has suggested that deregulation of P4HA2 is common in cancer. However, the expression pattern and molecular mechanisms of P4HA2 in glioma remain unknown. Here, we demonstrate that P4HA2 is overexpressed in glioma and inversely correlates with patient survival. Knockdown of P4HA2 inhibits proliferation, migration, invasion, and epithelial-to-mesenchymal transition (EMT)-like phenotype of glioma cells *in vitro* and suppressed tumor xenograft growth *in vivo*. Mechanistically, bioinformatics analysis shows that ECM-receptor interaction and PI3K/AKT pathway are the most enriched pathways of the co-expressed genes with P4HA2. Furthermore, P4HA2 mRNA was positively correlated with mRNA expressions of a series of collagen genes, but not mRNA of PI3K or AKT1/2. Conversely, both the protein expressions of collagens and phosphorylated PI3K/AKT could be downregulated either by silencing of P4HA2 expression or inhibition of its prolyl hydroxylase. Moreover, the inhibitory effects on the migration, invasion and the EMT-related molecules by P4HA2 knockdown can be recapitulated by the Akt phosphorylation activator. Taken together, our findings for the first time reveal an oncogenic role of P4HA2 in the glioma malignancy. By regulating the expression of fibrillar collagens and the downstream PI3K/AKT signaling pathway, it may serve as a potential anti-cancer target for the treatment of glioma.

**Highlights:** P4HA2 is overexpressed and correlated with poor prognosis in glioma.

P4HA2 depletion inhibits glioma proliferation, migration, invasion and EMT-like phenotype in vitro and tumorigenesis in vivo.

P4HA2 depletion attenuates the PI3K/AKT signaling pathway in a collagen-dependent manner.

## Introduction

Glioma is the most common adult malignant tumors in the central neural system, which is classified into different subtypes, with the glioblastoma multiforme (GBM) being the most abundant and malignant [1]. Despite the standard therapeutics including surgical resection, adjuvant radiotherapy and chemotherapy, the prognosis remains poorly improved [2]. Hence, great challenges still exist in the clarification of the molecular mechanism of glioma malignancy, which prompts us to identify novel molecular therapeutic target for GBM.

Recent years have witnessed increasing focus on the tumor microenvironment (TME) in the development of many solid tumors including GBM [3]. As one of the most robust features of TME, hypoxia can remodel the extracellular matrix (ECM), which may correlates with reprogramming of tumor cells and leads to tumor proliferation, invasion, epithelial-to-mesenchymal transition (EMT) and therapeutic resistance [4]. These proto-oncogenic effects of hypoxia are largely dependent on the activation and stabilization of hypoxia inducible factor (HIF), which mediates signaling transduction by transactivating a series of downstream target molecules [4]. Thus, the HIF targets deserve intense attentions for their pivotal roles in the hypoxia induced glioma malignancy.

Prolyl-4-hydroxylase subunit 2 (P4HA2) has been mentioned to be directly regulated by HIF in fibroblast and breast cancer, which can modulate the ECM in the TME [5]. In the previous work, we have demonstrated by high throughput microarray that the transcriptional level of HIF was significantly up-regulated in glioblastoma cell lines induced by hypoxia. Moreover, the binding site for HIF1α, termed as HIF reactive element(HRE), has also been identified by means of bioinformatics analysis(Dada not shown). Given the close relationship between hypoxia and cancer, it is reasonable to expect the involvement of P4HA2 in the maligmancy of glioma. In fact, deregulation of P4HA2 has been implicated in various pathological processes like osteoarthritis [6], hepatic fibrosis [7], giant cell arteritis [8], nonsyndromic high myopia [9] as well as many tumors such as hepatocellular carcinoma [7], breast cancer [10], papillary thyroid cancer [11], and oral cavity squamous cell carcinoma [12]. However, little is known about its expression pattern, biological functions and the role it may play in the oncogenesis of glioma.

In this study, we demonstrate that P4HA2 is overexpressed and correlated with poor prognosis in glioma. Knockdown of P4HA2 inhibits the proliferation, migration, invasion and EMT-like phenotype of hypoxia-cultured glioma cells *in vitro* as well as tumor xenograft growth *in vivo*. Mechanistically, knockdown of P4HA2 leads to inhibition of the PI3K/AKT pathway in a collagen-dependent manner. Thus, P4HA2 may serve as a potential therapeutic target for GBM patients.

## Materials and methods

### Clinical samples

Fresh frozen human tissue samples of 58 WHO grades II, III, and IV primary gliomas and 5 non-neoplastic brain tissues from surgical procedures for epilepsy were obtained from General Hospital of Jinan Military Command. All procedures related to acquiring the samples from the patients were given consent by the patients and were approved by the ethics committee of General Hospital of Jinan Military Command.

### Cell culture and reagents

U251 and A172 cell lines were purchased from the Chinese Academy of Sciences Cell Bank. The U87MG and LN229 cell lines were gifts from the laboratories of Dr. Hai-Zhong Feng (Renji-Med X Clinical Stem Cell Research Center, Shanghai Jiao Tong University) and Dr. Fang Wu (Shanghai Center for Systems Biomedicine, Shanghai Jiao Tong University). The authenticity of all these GBM cell lines was tested by short tandem repeat profiling. All cell lines were cultured in DMEM medium supplemented with 10% (v/v) fetal bovine serum (FBS), 100 U/ml penicillin, and 100 U/ml streptomycin from Gibco (Carlsbad, CA, USA) at 37°C in a 5% CO_2_ atmosphere. The P4HA2 enzymatic inhibitor Ethyl 3,4-dihydroxybenzoate (DHB, E24859) and phosphorylated AKT agonist SC79 (S7863) were purchased from Sigma-Aldrich (St Louis, MO, USA) and Selleck Chemicals (Huston, TX, USA) respectively. *Lentivirus transfection.* Short hairpin RNA (shRNA) against human P4HA2 (sh-P4HA2-1: CCGG**GCCGAATTCTTCACCTCTATT**CTCGAG **AATAGAGGTGAA GAATTCGGC**TTTTG;sh-P4HA2-2:CCGG**GCAGTCTCTGAAAGAGTACAT**CT CGAG**TGTACTCTTTCAGAGACTGC**TTTTTG) or a non-targeting scramble shRNA(Shanghai GenePharma, China) were cloned into vector LV3 (pGLVH1/GFP+Puro) and transfected into 293T cells using Lipofectamine 3000 (Invitrogen, CA, USA) for the generation of lentiviral vectors. The complete lentiviruses containing either P4HA2-specific or scramble shRNA(control) were then used to infect U251 and U87MG cell lines and selected by puromycin for 48 h after infection.

### Proliferation assay

Cell proliferation was detected by WST-1 assay at 24, 48, 96 and 120 h according to the manufacturer’s instructions. Briefly, cells were seeded into 96-well plates at a density of 5×10^3^ cells in 200 μl medium per well. At indicated time point, wst-1 reagent (Roche, Basel, Switzerland) was added to each well and incubated for 1 h. The absorbance rate at 450 nm was read by using a microplate reader (Bio-Rad Laboratories, CA, USA.). All experiments were performed in triplicate and repeated at least three times.

### Invasion assay

*T*umor invasion was determined by Boyden Chamber assay (Corning, 3422, NY, USA) with BioCoat™ Matrigel™ (BD, 356234, NY, USA). Cells were serum-starved for 24 h; 5×10^4^ cells were plated into the upper well of the Boyden chambers with serum-free medium in the top chamber and medium with 10 % FBS in the lower chamber. After 24 h of incubation, cells on the top of the membrane were removed. Cells that invaded through to the bottom surface of the membrane were washed with PBS, fixed in 4 % paraformaldehyde and stained with 0.2 % crystal violet. The migrated cells were observed under Leica inverted microscope (Solms, Germany). Cell number was counted in eight random fields for each condition. The experiments were performed in triplicate.

### Migration assay

Tumor migration was determined by wound healing assay. Cells were seeded in 6-well plates and cultured to 80-90% confluence. After serum starvation for 12 h, a wound was created by scraping the cell monolayer with a 200μl pipette tip. The cultures were washed by PBS to remove the floating cells. Then the cells were cultured in serum-free medium. Cell migration into the wound was observed at the indicated times (0 h, 36 h) in marked microscopic fields and images were captured with a DS-5M Camera System (Nikon, Tokyo, Japan). The data obtained were presented as a migration rate by dividing the distances between wound edges with ImageJ software. *Quantitative RT-PCR analysis.* The procedure of qRT-PCR had been previously described [13]. The gene expression was calculated using the 2^-ΔΔ^ Ct method. All data represent the average of three replicates. The PCR primer sequences were listed as follows. P4HA2, forward 5’-CAAACTGGTGAAGCGGCTAAA-3’, reverse 5’-GCACAGAGAGGTTGGCGATA-3’; SNAI1, forward 5’-TCGGAAGCCTAACTACAGCGA-3’, reverse 5’-AGATGAGCATTGGCAGCGAG-3’; SLUG, forward 5’-CGAACTGGACACACATA CAGTG-3’, reverse 5’-CTGAGGATCTCTGGTTGTGGT-3’; TWIST1, forward 5’-GTCCGCAGTCT TACGAGGAG-3’, reverse 5’-GCTTGAGGGTCTGAATCTTGCT-3’; GAPDH, forward 5’-CACCCACTCCTCCACCTTTG-3’, reverse 5’-CCACCACCCTGTTGCTGTA G-3’

### Western blot

The standard protocol of western blot had been described previously [14]. In brief, the total cell lysates were prepared in high KCl lysis buffer (10 mmol/L Tris-HCl, pH 8.0, 140 mmol/L NaCl, 300 mmol/L KCl, 1 mmol/L EDTA, 0.5% Triton X-100, and 0.5% sodium deoxycholate) with complete protease inhibitor cocktail (Roche). The protein concentration was determined using a BCA Protein Assay Kit (Pierce, Rockford, IL, USA). Equal amounts of protein samples were subjected to SDS-PAGE and transferred to polyvinylidene fluoride (PVDF) membranes (Millipore, MA, USA). The membranes were treated with 1% blocking solution in TBS for 1 h, and immunoblots were probed with the indicated primary antibodies at 4°C overnight. The primary antibodies used were P4HA2 (ab233197), Collagen I (ab34710), Collagen III (ab7778) and Collagen IV (ab6586) from Abcam (Cambridge, UK), Twist1 (25465-1-AP) from Proteintech Group (Rosemont, PA, USA), Snail (#3879), Slug (#9585), E-cadherin (#3195), glyceraldehyde 3-phosphate dehydrogenase (GAPDH) (sc-47724) from Santa Cruz Biotechnology (Dallas, TX, USA). Then the membranes were washed and incubated with HRP-labelled secondary antibodies (Molecular Probes, Waltham, MA, USA). The fluorescence signals were detected by a BM Chemiluminescence Western Blotting kit (Roche, Basel, Switzerland). Densitometry quantification was calculated and analyzed using ImageJ software.

### Immunohistochemistry (IHC) staining

Tumors were fixed with 10% formalin, followed by paraffin embedding and sectioning. The protocol has been described in the previous study [14]. The percentages of positive tumor cells scored 0 (0% to 25%), 1 (26% to 50%), 2 (51% to 75%), and 3 (76% to 100%). The staining intensities were graded as follows: 0 (negative), 1 (weak), 2 (moderate), or 3 (strong). The final scores were the arithmetic product of the percentage and staining intensity, that is, (None) =0, (Low) = 1 to 3, (Medium) = 4 to 6, (High) = 7 to 9.

### Animal Studies

The procedures of animal studies had been mentioned previously [14]. All mouse experiments were carried out in accordance with institutional guidelines and regulations of the government. All mouse experiments were also approved by the Ethical Board at the General Hospital of Jinan Military Command.

### Immunofluorescence and hematoxylin and eosin (H&E) staining

The procedures of immunofluorescence and H&E staining had been previously described [14]. In brief, cells on poly-L-lysine-coated glass coverslips were fixed with 4% paraformaldehyde for 15 minutes at room temperature and permeabilized by ice-cold methanol for 10 minutes. The xenografts were fixed in 4% paraformaldehyde, embedded in optimal cutting temperature compound for freezing, and then cryosectioned (10-mm sections). After being blocked by 15% normal donkey serum for 30 minutes, the cells were incubated at room temperature for one hour with primary antibody diluted in antibody buffer. After incubation with the Ki-67 primary antibodies (Santa Cuz, sc-56319, Dallas, TX, USA), the cells were rinsed and incubated for one hour at room temperature with Alexa Fluor-labeled secondary antibodies (Molecular Probes, Waltham, MA, USA). The cells were washed with PBS and the cover slips were mounted with glycerine/PBS containing 0.1 mg/mL DAPI for nuclei staining. For H&E staining, the fixed tumor was embedded in paraffin, cut into 6 mm sections and stained with H&E. Slides were photographed using an optical or confocal microscope (Olympus, Tokyo, Japan).

### Bioinformatics

The data including P4HA2 transcriptional expression, Kaplan-Meier analysis of P4HA2 transcriptional level over the overall survival time as well as the correlation of P4HA2 transcription with other genes were downloaded from the TCGA dataset including both GBM and LGG samples(The Cancer Genome Atlas). Data visualization was performed on the web server GEPIA (http://gepia.cancer-pku.cn) [15,16]. According to the transcriptional level of P4HA2 derived from TCGA, glioma samples were divided into two subgroups with higher or lower P4HA2 expression respectively. Differentially expressed genes(DEGs) of the two subgroups were analyzed using the Genespring GX software (Agilent Technologies, USA). Genes with an absolute fold-change ⩾ 2 and an adjusted p-value (FDR) < 0.05 in expression were considered as DEGs and were subsequently mapped onto GO and KEGG using DAVID tool (http://david.abcc.ncifcrf.gov/).

### Statistical analysis

All experiments were performed in triplicates and repeated at least thrice unless indicated otherwise. Statistical analysis was performed in GraphPad Prism software (Version 7, CA, USA). Experimental data were expressed as means ± standard deviation (SD). IHC staining scores were analyzed by Mann Whitney test for two groups and Kruskal-Wallis test for multiple comparisons respectively. Continuous data were calculated with Student’s t test for two group comparison and AVONA for multiple comparisons. Overall survival of patients with different P4HA2 protein levels was compared with Kaplan–Meier and log-rank test. The transcriptional correlations were analyzed using Pearson correlation coefficient. P values less than 0.05 were considered as significant.

## Results

### P4HA2 is overexpressed and correlated with poor prognosis in glioma

To examine the role of P4HA2 in glioma, we first compared the tumor expression of P4HA2 with normal tissues at the transcriptional level. By interrogating the TCGA, we found the mean transcriptional level of P4HA2 in glioma samples was higher than that in normal tissues (Fig. 1A, p<0.05). We next examined the relationship between P4HA2 transcriptional level and histologic grade in human glioma. As expected, we found increased P4HA2 transcriptional level was associated with increasing histologic tumor grade in glioma, which was mostly elevated in WHO IV glioma (Fig. 1B, p<0.001). Then, we took advantage of the survival data in TCGA to investigate the correlation between P4HA2 transcriptional expression and prognosis of patients. The results revealed that the overall survival of patients with higher P4HA2 transcriptional expression was notably shorter than that of patients who have lower P4HA2 expression in GBM(Fig. 1C, p<0.001). Meanwhile, immunohistochemistry in clinical samples, showed that P4HA2 was extensively expressed in the cytoplasm and the median P4HA2 post-translational level in 58 glioma samples was significantly higher than that in 5 normal tissues. Similar to the mRNA expression, we found the expressional level of P4HA2 protein increased along with the increased pathological grade of tumors (Fig. 1D-F). At last, by log-rank test, the overall survival of patients with higher post-translational P4HA2 level was found to be shorter than those expressing lower P4HA2 protein, which was consistent with the transcriptional results, implying that either mRNA or protein expression of P4HA2 was negatively associated with the prognosis (Fig. 1G, p<0.05).

**Figure 1.**
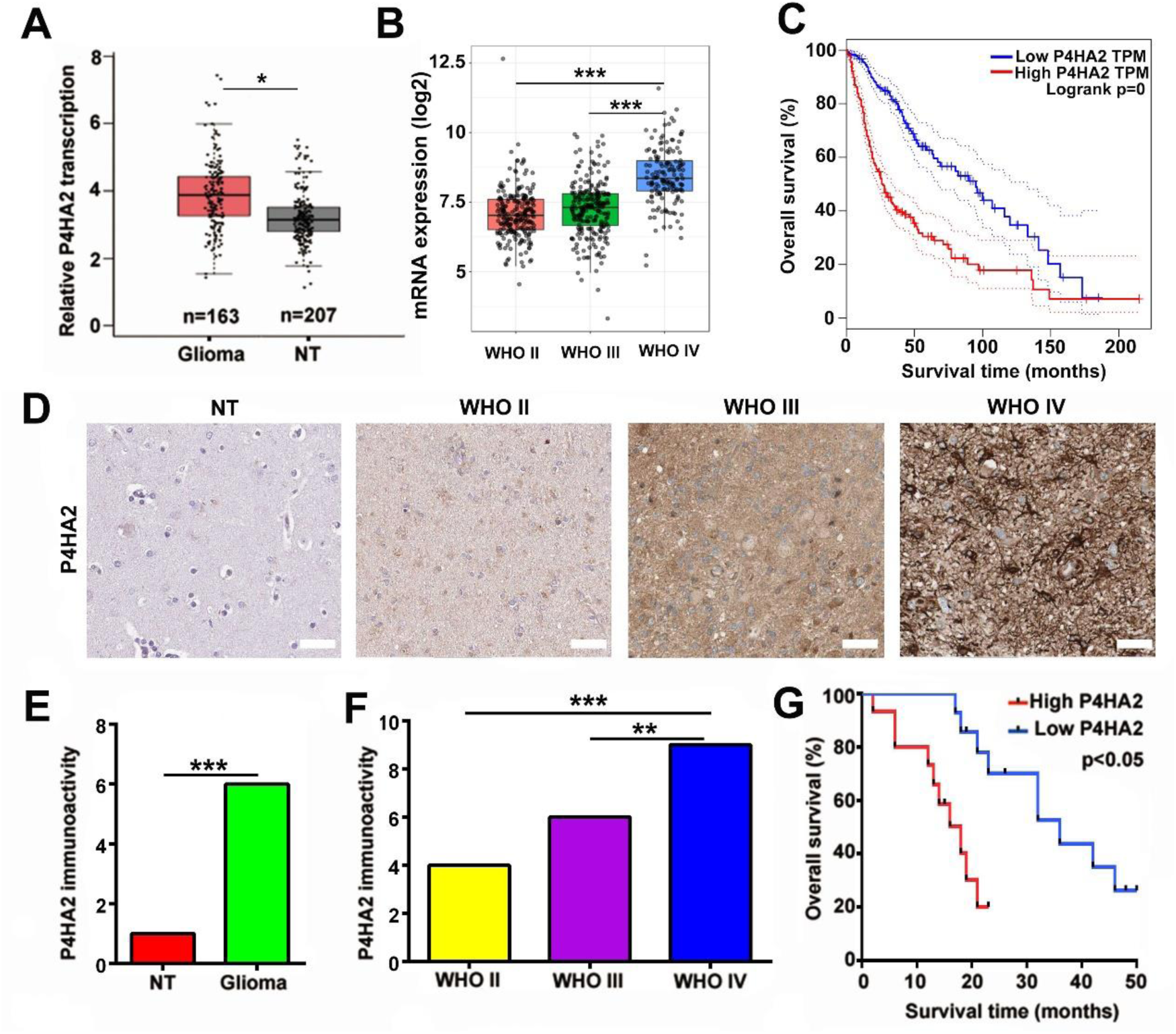
P4HA2 is overexpressed and correlated with poor prognosis in glioma. (A) Statistical comparison of P4HA2 transcriptional expression between normal tissues and glioma tissues in the TCGA database. (B) Comparison of P4HA2 mRNA expression between glioma with different pathological grades(WHO II, III, IV) from TCGA; ***p<0.001. (C) Kaplan–Meier result showed the overall survival discrepancy between the glioma patients with high and low P4HA2 transcriptional expressions from TCGA. (D) Representative images of cytoplasmic P4HA2 expression in normal brain tissues and glioma samples by IHC staining; scale bar=50 μm. (E) Comparison of P4HA2 IHC expression between normal and glioma tissues and (F)between glioma samples with different pathological grades; ***p<0.001. (G) Kaplan–Meier analysis showing the overall survival discrepancy between patients with high and low P4HA2 IHC expressions; p<0.05.

### Knockdown of P4HA2 inhibits proliferation, migration and invasion of glioma cells in vitro and suppresses tumorigenesis in vivo

To explore the role of P4HA2 in glioma malignancy, the U251 and U87MG cells were chosen for the RNA interference experiment because of their relatively high P4HA2 expression level among the four GBM cell lines (Fig. 2A). It can be seen that both shRNAs targeting P4HA2 mRNA were effective to inhibit the expression of P4HA2 (Fig. 2B). Malignant proliferation was among the most distinct features of glioma (ie. GBM). In view of this, WST-1 assay was performed to determine the impact of P4HA2 on cell proliferation. After stable P4HA2 knockdown, both U251 and U87MG cells showed obviously reduced rate of proliferation compared to the control group at each time point(Fig. 2C, p<0.05). Notably, glioma cells transfected with sh-P4HA2-2 lentivirus had a relatively lower proliferation rate than cells with sh-P4HA2-1 transfection, thus being selected for the further experiments. To clarify the role of P4HA2 interference in cell invasion, which is another aspect of glioma malignance, Matrigel invasion assay was performed to reveal that P4HA2 knockdown significantly decreased the number of invading cells in both U251 and U87MG cells at 24 hours after serum-free culture (Fig. 2C, p<0.001). Moreover, wound-healing assay showed that the migration rate of cells in both glioma cell lines were markedly reduced at 36 hours (Fig. 2D, p<0.001).

**Figure 2.**
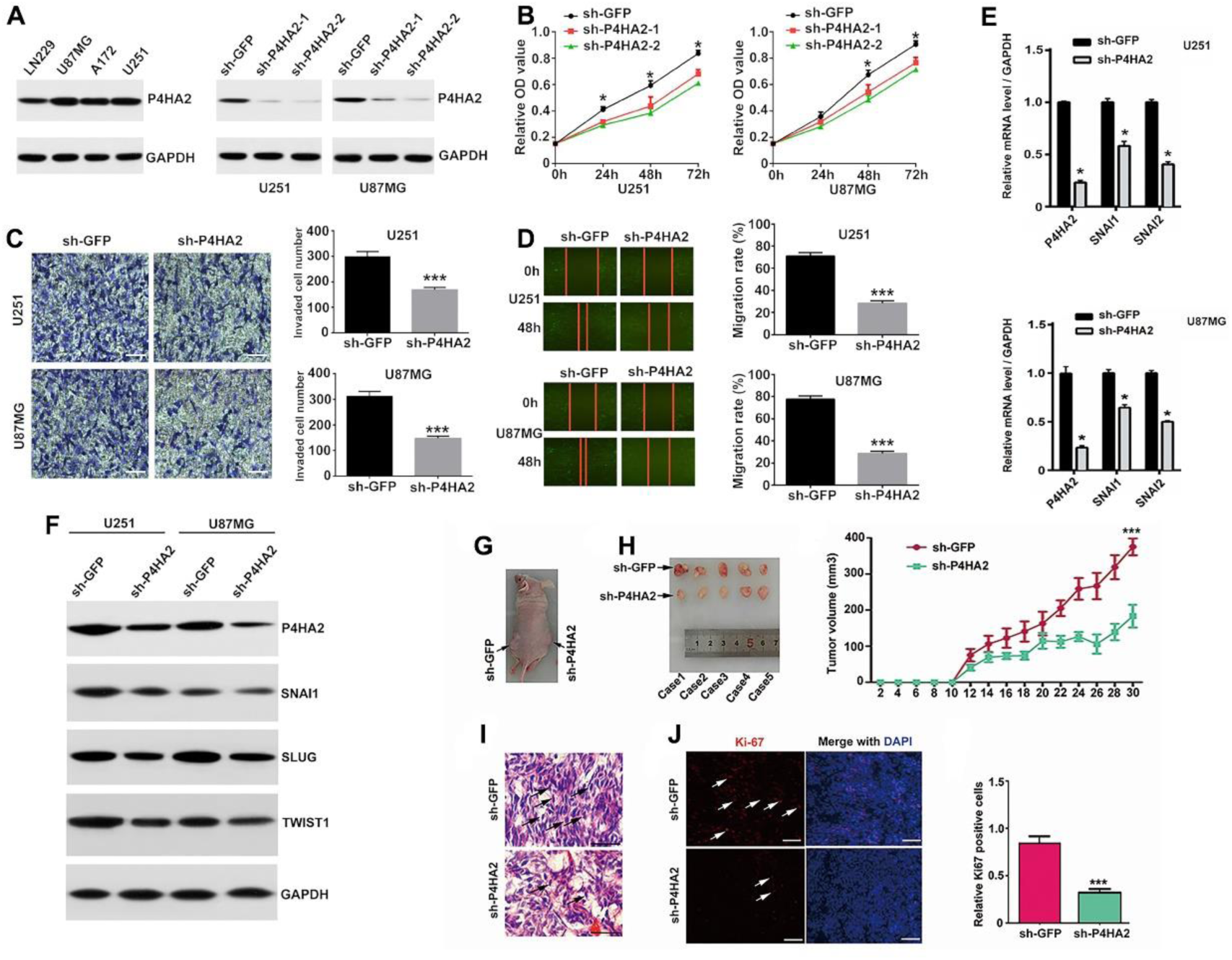
Knockdown of P4HA2 inhibits malignant biological behavior of glioma cells *in vitro* and *vivo*. (A) P4HA2 expression level was detected by western blot in several established GBM cell lines(left). The efficiency of two P4HA2 shRNAs were validated by western blot in U251 and U87MG cells(right). (B) Cell proliferation was detected by wst-1 assay at 0, 24, 48, 72h in U251 and U87MG cells(*p<0.05). (C) Cell invasion of both cell lines was assessed in BioCoat™ Matrigel™ Invasion Chamber by crystal violet staining(left, Scale bar=100μm); invaded cell number was shown in histogram on the right(***p<0.001). (D) Cell migration was observed and photographed at 48h(left). Data were presented as percentage of gap distance at 0h compared to 48h and shown in histograms(right, ***p<0.001). (E) qRT-PCR and (F)immunblotting results in U251 and U87MG cell lines showed downregulation of EMT-promoting molecules after shP4HA2 transfection. (G) Nude mice were subcutaneously implanted with P4HA2 control and P4HA2 knockdown U87MG cells on either side to generate xenograft tumors. (H) Photographs represented removed tumor bulks from two groups of animals with subcutaneous xenografts(left). Tumor volume from mice implanted with P4HA2 knockdown U87MG cells were compared with the control tumor at indicated days(right,***P < 0.001). (I) H&E staining of xenograft tumor tissues. Bar=25μm. (J) Immunofluorescence images showing expression levels of Ki-67 (red) in P4HA2 control tumors and P4HA2 knockdown tumors. DAPI (blue) was used to stain nuclei(left, Scale bar=100μm). Statistical comparison of the relative Ki-67 expression between groups was shown on the right(***P < 0.001).

Subsequently, we tried to further verify the effect of P4HA2 on tumorigenesis *in vivo*, where the ECM may play a more important role than *in vitro*. Through subcutaneous xenograft assay in a nude mouse model with U87MG cells, which was the mostly inhibited cell line in the *in vitro* experiment, we found that knockdown of P4HA2 significantly inhibited tumor growth in the observed post-implantation period(Fig. 2F-G). What is more, tissue H&E staining showed that the tumor cells with characteristics of abnormal morphology were obviously decreased in the xenograft derived from P4HA2-knockdown cells (Fig. 2I). Consistently, fluorescent staining of cell proliferation marker Ki67 merged with nucleus showed decreased ratio of Ki-67 immunoactivity in P4HA2-knockdown xenograft compared to the control xenograft (Fig. 2J, K).

### Knockdown of P4HA2 inhibits the EMT-like phenotype in glioma cells

Stemeness and EMT are both hallmarks of malignant tumor cells which intensively participate in cancer invasion and metastasis. By EMT transition to stromal cells, tumor cells gain misplaced stem-like properties, including the ability of aberrantly communicating with peri-tumor environment(ie. ECM) for invasive growth into adjacent tissues[16]. Thus it’s reasonable to obtain insight into the role of P4HA2 in the regulation of EMT of glioma cells. We first examined the relation between P4HA2 and EMT expression using TCGA database, and interestingly found a positive correlation of P4HA2 mRNA expression with a series of EMT-promoting markers (Fig. S1). Beyond this, quantitative analysis by qRT-PCR exhibited more than 2-fold reduction in the mRNA expression of SNAI1, SLUG, and TWIST1 in both glioma cell lines. Similarly, the protein expressions of these molecules were notably reduced which were seen by qualitative immunoblot assay. The results suggest that the expression of SNAI1, SLUG, and TWIST1 are regulated by P4HA2 either in transcriptional or translational level. Taken together, the data presented in Figure 2 demonstrate that P4HA2 promotes proliferation, invasion, migration and EMT phenotype as well as *in vivo* tumorigenesis of glioma cells.

### P4HA2 regulates collagen deposition in glioma

Subsequently, we made further attempt to delineate the molecular underpinning by refining the gene expression profiles of glioma patients from TCGA database. These patient data were grouped on the basis of their P4HA2 expressional levels, and the differentially expressed genes(DEGs) between the two P4HA2 expressional groups were compared by KEGG pathway analysis. A slew of pathways tightly correlated with P4HA2 expression were exhibited in Figure 3A. Of particular interest was the finding that ECM-receptor interaction bears the most enrichment of the DEGs. Abnormal remodeling of ECM are characteristic of tumor microenvironment. As the primary components of ECM fibers, collagens intensively involved in the adhesion, elongation and motility of tumor cells, and collagen deposition within tumor cells has been proved to promote tumor invasion and metastasis. As the substrate of collagen hydroxylase, are collagen contents in the glioma sample also regulated by P4HA2? First of all, we found by TCGA database the expressional profiles of a spectrum of collagens in high-P4HA2 group were increased at least 2-fold compared to low-P4HA2 group(Fig.S2). There were significantly positive correlation in between P4HA2 mRNA and most of those collagen genes(Fig.3B). Next, the knockdown experiment by immunoblotting in glioma cell lines intriguingly confirmed that collagen I, III and IV, three representative collagens in the brain, are in regulation of P4HA2 not only in the content of mRNA but also protein(Fig.3C).

**Figure 3.**
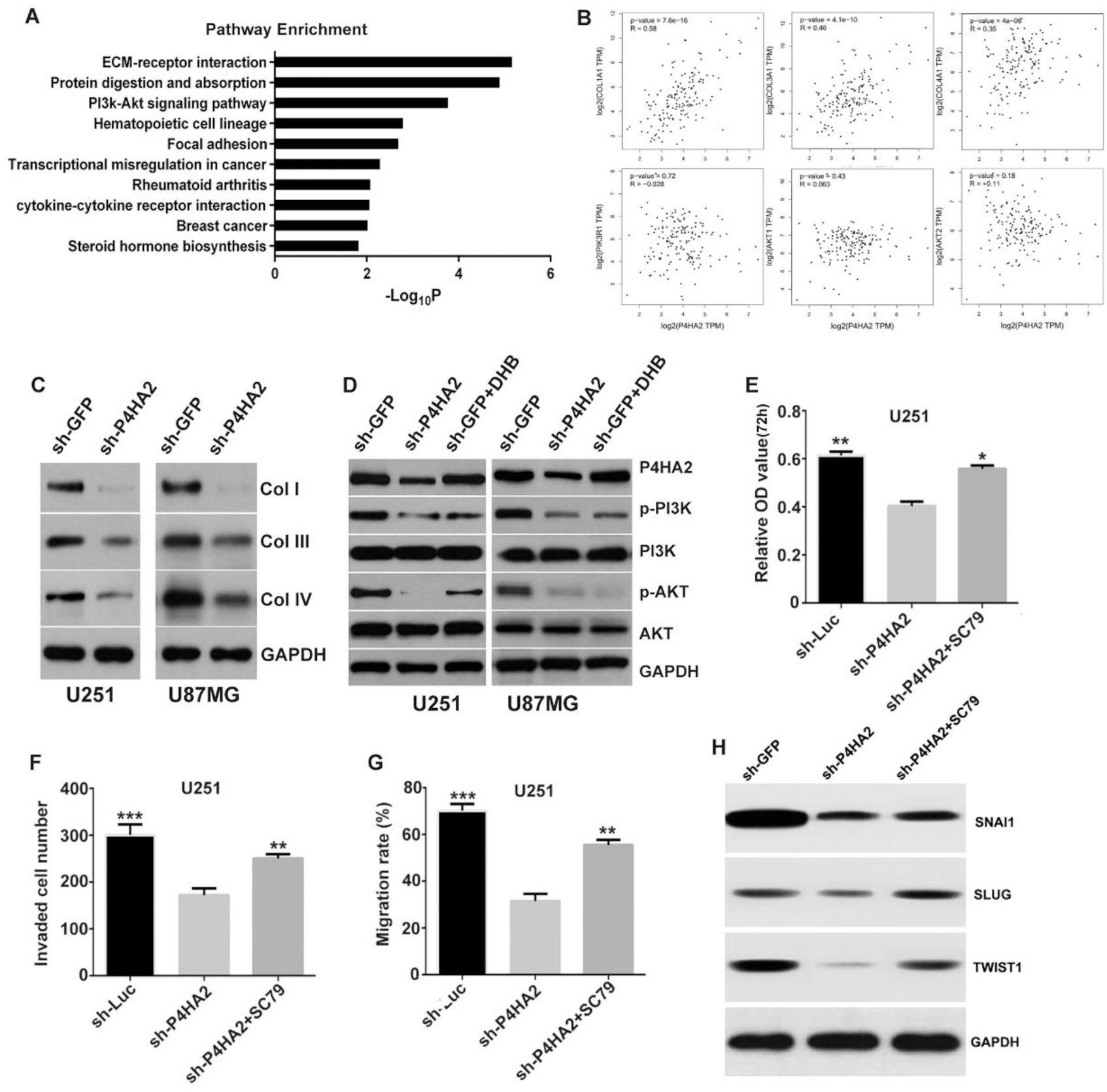
Knockdown of P4HA2 attenuates the collagen-dependent PI3K/AKT signaling pathway. (A) KEGG pathway analysis based on TCGA database figured out the enriched pathways associated with P4HA2 expression. (B)Transcriptional correlations of P4HA2 with collagens(upper) and PI3K/AKT signaling molecules(lower) were exhibited with TCGA analysis. (C) Immunoblot assay showed the regulation of three representative collagen types I, III and IV by P4HA2 knockdown. (D) The effect of P4HA2 depletion or enzyme inhibition on the PI3K/AKT signaling pathway was determined by immunobolt assay in two glioma cell lines. (E) The inhibitory effect on glioma proliferation, (F)cell invasion and (G)migration by P4HA2 depletion was recapitulated by AKT phosphorylation activator SC79; *p<0.05, **p<0.01, ***p<0.001. (H) Immunobolt assay also showed rescued expression of previously downregulated EMT related factors by P4HA2 knockdown.

### P4HA2 regulates PI3K/AKT pathway of glioma cells in a collagen dependent manner

Previous studies have determined that PI3K/AKT signaling is among the most critical pathways in the irregular molecular networks underlying the tumoregenesis of glioma[17]. In the current study bioinformatics was utilized to show that PI3K/AKT is also among the most enriched pathways associated with P4HA2 expression[17, 18]. We proposed a hypothesis that P4HA2 is required for the activation of PI3K/AKT pathway. Interestingly, though the mRNA levels PI3K and AKT were not influenced by P4HA2 knockdown(Fig.3B), we found by immunoblotting that P4HA2 inhibition led to reduced protein levels of phosphorylated PI3K and AKT. To our knowledge, collagens can transduce intracellular signaling through the receptor tyrosine kinases(RTKs) on the cell membrane surface[19]. To test whether it is collagen that regulates the PI3K/AKT pathway, we utilized a P4HA2 enzyme inhibitor, DHB to suppress the deposition of collagen in glioma cells. The results were the same: both phosphorylated PI3K and AKT expressions, not the total protein expressions, were significantly reduced by inhibition of the collagen hydroxylase, implying that P4HA2 regulates PI3K/AKT signaling in a collagen-dependent manner(Fig.3D). At this point, we were strongly inspired to find out whether PI3K/AKT activation is able to mediate the pro-tumoregenesis effect of P4HA2. By employing an AKT agonist SC79, the inhibited proliferation, invasion and migration in the P4HA2 knockdown cell line were all shown to be notably reversed (Fig. 4E-G). Specially, the EMT related molecules, which have been downregulated by P4HA2 knockdown, were also found to increase after AKT activation, which confirms and extends the previous finding that EMT is tightly associated with AKT family(Fig. 4H). Taken together, this part of results indicates the transduction of collagen-PI3K/AKT, which is under the regulation of P4HA2, is possibly the underlying mechanism for the tumor-promoting effect of P4HA2.

**Figure 4.**
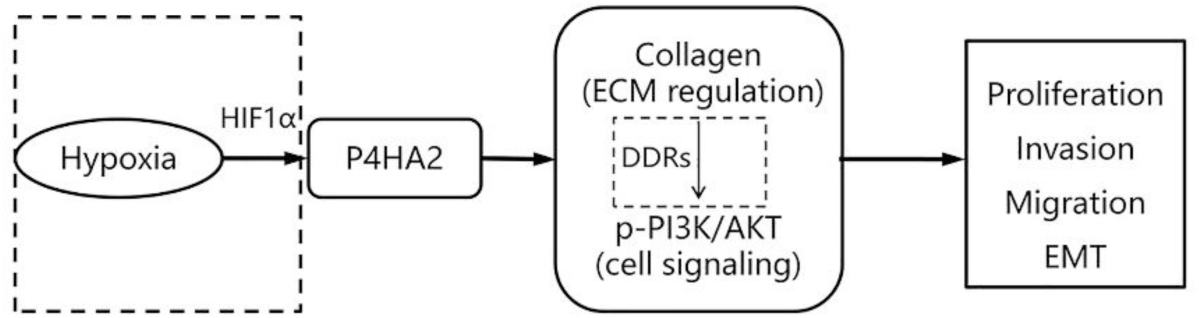
A working model of the mechanism for the pro-tumorigenesis of P4HA2. HIF1α induced P4HA2 promotes the malignances of glioma cells by regulating collagen deposition and the downstream PI3K/AKT pathway.

## Discussions

In this study, we firstly investigated the expression pattern and clinical significance of P4HA2 in glioma patients. We found that P4HA2 was frequently upregulated in glioma, which was consistent with its expression in other malignancies [7, 10-12]. The inverse correlation between P4HA2 and patient survival indicated that P4HA2 may serve as a promising prognostic marker in glioma. Then, the biological functions of P4HA2 was examined. Similar with the findings in breast cancer [20], knockdown of P4HA2 was also found to inhibit glioma proliferation, migration and invasion *in vitro* and tumor xenograft growth *in vivo.* However, the underlying molecular mechanism is elusive.

EMT, a biological process characterized by the downregulation of epithelial markers represented by E-cadherin, and the upregulation of mesenchymal markers like vimentin, fibronectin, and several key EMT inducing factors including snail, slug and twist, is a pivotal step for cell migration and invasion in various cancer types including glioma [21, 22]. Hence, the relation between P4HA2 and EMT was examined. By bioinformatic analysis, we showed that P4HA2 expression was correlated with a series of EMT-associated factors in glioma patients. Knockdown of P4HA2 was accompanied by elevation of E-cadherin and reduction of vimentin, fibronectin, snail, slug and twist, which indicated that the EMT-like phenotype may participate in the P4HA2-mediated glioma migration and invasion. However, the mechanism for P4HA2-induced cell proliferation and EMT-like phenotype still needs to be further elucidated.

By a preliminary investigation through pathway analysis of the difference between the gene expression profiles of high- and low-P4HA2 glioma patients from TCGA database, we found that the most enriched pathway was ECM-receptor interaction, which implicated that the mechanism of P4HA2 may lie in the interaction between ECM and glioma cells. Among these pathways, the PI3K/AKT signaling pathway attracted our attentions. Multiple studies have shown that the PI3K/AKT signaling pathway is a crucial cascade for cancer proliferation, migration, invasion, metastasis and therapeutic resistance [23, 24]. Besides, PI3K/AKT signaling pathway is also a common upstream regulator of the EMT-like phenotype [25]. Hence, the PI3K/AKT signaling pathway is considered as a probable mechanism for P4HA2 in glioma. By western blot, we found the PI3K/AKT signaling pathway could be either inactivated by silencing the expression or blocking the enzyme activity of P4HA2. This may indicate that the regulation of PI3K/AKT signaling pathway by P4HA2 is in a collagen-dependent manner. Then, through rescue assay with a signal agonist SC79, we further confirmed that the PI3K/AKT signaling pathway was factually required for the oncogenic role of P4HA2. Thus, the PI3K/AKT signaling pathway is considered to account for the effects of P4HA2 on glioma proliferation, migration, invasion and EMT-like phenotype. However, how to link collagen with PI3K/AKT signaling pathway needs further clarification.

As the major frame component of ECM, a series of collagens including types I, III, IV, VI and XVI have been mentioned to be upregulated in glioma [26-30], which demonstrates a crucial role of collagen in gliomagenesis. In brain tumors, collagen has three primary functions. Apart from acting as a scaffold for cell adhesion and serving as a reservoir of matricellular molecules, collagen can act as a ligand for ECM-cell signalling transduction through interacting with binding receptors [31, 32]. Currently, three types of collagen receptors have been identified in GBM. The first is integrins, a class of transmembrane receptors, which can mediate signal transduction through formation of a large adhesion complex containing signal molecules [33]. The second is discoidin domain receptors (DDRs), a class of receptor tyrosine kinases, which can mediate glioma progression via activation of downstream signal molecules [34]. The third is Endo 180, a member of the mannose receptor family, which can mediate collagen internalization and glioma invasion [35]. Among the collagen receptors, the DDRs has been tightly associated with the PI3K/AKT signaling pathway, because of the C-terminal of DDR1 kinase domain contains the YELM binding motif for association with the p85 subunit of PI3K [36]. As previous literature has demonstrated that type I-IV collagens can all combine with DDRs [37, 38] and deposited by P4HA2 [6, 20], we think at least the DDRs could theoretically mediate the collagen-dependent regulation of PI3K/AKT signaling pathway by P4HA2 in glioma, though it needs further confirmation. With the downregulation of collagen I, III and VI by P4HA2 silencing in glioma cells, a novel signal axis of P4HA2-collagen-PI3K/AKT pathway is proposed in contribution to glioma progression. A working model of P4HA2 was proposed whereby P4HA2 catalyzes collagen deposition, promotes the interaction between collagens and their receptors to activate the PI3K/AKT signaling pathway, which leads to glioma proliferation, migration, invasion and the EMT-like phenotype However, the activation of PI3K/AKT axis by SC79 is insufficient to rescue the inhibitory effects of P4HA2 depletion, which indicates that other mechanisms may also exist. For example, previous study has mentioned that the ECM rigidity can regulate the proliferation, mobility and EMT of glioma cells[39, 40]. Whether P4HA2 can promote glioma progression by directly enhancing ECM rigidity(changing the physical property of ECM) through depositing collagens needs further exploration. Besides, considering the focal adhesion kinase (FAK) signaling pathway is another oncogenic mechanism for glioma malignant behaviors [41], the collagen-dependent regulation of focal adhesion by P4HA2 also merits attention. Moreover, P4HA2 may also enhance glioma stenmess and neovascularization, because the same function has been reported in P4HA1, an isoform of P4HA2, by promoting glioma stem cell-endothelial cell transdifferentiation and the collagen IV dependent formation of vascular basement membrane [42]. Besides collagen-dependent mechanism, P4HA2 has recently been reported to promote B-cell lymphoma progression via hydroxylation of Carabin [43]. This indicates that the substrates of P4HA2 are not limited to collagens, which is beneficial for a deeper understanding of the role played by P4HA2 in cancer biology. In conclusion, our study first demonstrates that P4HA2 is a potential oncoprotein participating in glioma pathogenesis, which may serve as a prognostic marker in glioma. Besides, we reveal a potential P4HA2-collagen-PI3K/AKT pathway in mediating the glioma progression, which sheds light on the importance of interplay between glioma cells and their microenvironment in glioma progression. Targeting P4HA2 by developing specific enzymatic inhibitors or RNA interference may represent a potential therapeutic strategy in the treatment of GBM.

## Acknowledgements

This research was supported by the National Natural Science Foundation of China (81702944, 81402063 and 81602193), Medical Research Youth Innovation Fund of Sichuan Province(Q18035) and Shanghai Jiao Tong University Med-X Fund (YG2015MS20). We thank Dr. Hai-Zhong Feng and Dr. Fang Wu for their gift of glioma cell lines.

